# Arc ubiquitination regulates endoplasmic reticulum-mediated Ca^2+^ release and CaMKII signaling

**DOI:** 10.1101/2022.09.02.506383

**Authors:** Mohammad A. Ghane, Zachary D. Allen, Cassandra L. Miller, Bin Dong, Jenny J. Yang, Ning Fang, Angela M. Mabb

**Affiliations:** Georgia State University, Neuroscience Institute, Atlanta, GA USA; Georgia State University, Center for Behavioral Neuroscience, Atlanta, GA USA; Georgia State University, Department of Chemistry, Atlanta, GA USA

**Keywords:** Arc, CaMKII, endoplasmic reticulum, calcium, ubiquitination

## Abstract

Synaptic plasticity relies on rapid, yet spatially precise signaling to alter synaptic strength. Arc is a brain enriched protein that is rapidly expressed during learning-related behaviors and is essential for regulating metabotropic glutamate receptor-mediated long-term depression (mGluR-LTD). We previously showed that disrupting the ubiquitination capacity of Arc enhances mGluR-LTD; however, the mechanism by which this occurs and its consequences on other mGluR-mediated signaling events is unknown. Here we show that disrupting Arc ubiquitination on key amino acid residues leads to derangements in Ca^2+^ release from the endoplasmic reticulum (ER) during pharmacological activation of Group I mGluRs. These alterations were observed in all neuronal subregions except secondary branchpoints. Deficits in Arc ubiquitination increased Arc self-association and enhanced its interaction with calcium/calmodulin-dependent protein kinase IIb (CaMKIIb) and constitutively active forms of CaMKII. Notably, these interactions were also excluded at secondary branchpoints. Finally, disruptions in Arc ubiquitination were found to increase Arc interaction with the integral ER protein Calnexin. These results suggest a previously unknown role for Arc ubiquitination in the fine tuning of ER-mediated Ca^2+^ signaling that is needed for mGluR-LTD, which in turn, may regulate CaMKII and its interactions with Arc.

## 1 Introduction

Synaptic plasticity relies on rapid, spatially precise signaling to appropriately alter synaptic strength. mGluR-LTD is a protein synthesis-dependent form of plasticity that is initiated by activation of G_q_-coupled receptor pathways, activating ER-bound inositol triphosphate receptors (IP_3_Rs) to allow Ca^2+^ efflux, which is canonically associated with α-amino-3-hydroxy-5-methyl-4-isoxazolepropionic acid (AMPA) receptor endocytosis (Abe et al., 1992; Aramori and Nakanishi, 1992; Palmer et al., 1997; Snyder et al., 2001; Wilkerson et al., 2018). It is established that neurons contain highly complex continuous ER networks, with ER morphology increasing in complexity at dendritic branchpoints and [Ca^2+^]_ER_ varying at different compartments (Terasaki et al., 1994; Spacek and Harris, 1997; Cui-Wang et al., 2012; Krijnse-Locker et al., 2017). Furthermore, the distribution of IP_3_Rs vary throughout the neuron, with higher levels found at proximal regions (Sharp et al., 1993; Blaustein and Golovina, 2001). It is hypothesized that these various gradients offer an additional layer of local protein synthesis and trafficking regulation.

Activity-regulated cytoskeleton-associated protein (Arc) is vital for numerous forms of synaptic plasticity, including mGluR-LTD (Lyford et al., 1995; Chowdhury et al., 2006; Plath et al., 2006; Shepherd et al., 2006; Beique et al., 2011; Korb and Finkbeiner, 2011; Jenks et al., 2017; Pastuzyn and Shepherd, 2017; Mabb and Ehlers, 2018; Newpher et al., 2018). Arc induction is spurred by learning and memory related behaviors, including spatial memory acquisition, consolidation, and retrieval (Guzowski et al., 1999; Ramirez-Amaya et al., 2005; Plath et al., 2006; Bramham et al., 2008; Nakayama et al., 2015). Importantly, Arc is rapidly degraded largely through proteasome- and lysosome-dependent processes (Rao et al., 2006; Yan et al., 2018). Proteasome-dependent Arc degradation is facilitated by ubiquitination of two tandem lysine residues (268/269) catalyzed by the E3 ubiquitin ligase RNF216 (Mabb et al., 2014). We previously generated a transgenic mouse that contained mutations in the Lysine sites 268/269 to Arginine (ArcKR), which resulted in Arc build up while retaining endogenous control of its expression. ArcKR mice have alterations in AMPA receptor trafficking, enhanced mGluR-LTD, and deficits in spatial reversal learning, including a bias in utilizing inefficient search strategies (Wall et al., 2018).

Calcium/calmodulin-dependent protein kinase II (CaMKII) is another hub of synaptic plasticity known to be essential for learning (Coultrap et al., 2014; Murakoshi et al., 2017; Rossetti et al., 2017; Bayer and Schulman, 2019). CaMKII activity is tightly regulated by extra- and intracellular Ca^2+^ dynamics, resulting in autophosphorylation at specific threonine residues that determine activity states (Glazewski et al., 2000; Elgersma et al., 2002; Gillespie and Hodge, 2013). Previous work has shown that CaMKII plays a role in mGluR-LTD (Mockett et al., 2011; Cook et al., 2022) and that dysregulation of CaMKII leads to LTD enhancement and alterations in spatial search strategies (Bach et al., 1995). Arc interacts with CaMKII, which is believed to recruit Arc to different subdomains (Donai et al., 2003; Okuno et al., 2012). Recent evidence indicates that CaMKII phosphorylates the GAG domain of Arc, promoting Arc oligomerization and increasing mGluR-LTD magnitude (Zhang et al., 2019; Zhang and Bramham, 2020).

Given the importance of Arc mRNA and protein localization to specific neuronal subregions (Steward et al., 1998; Moga et al., 2004; Chowdhury et al., 2006; Peebles et al., 2010; Okuno et al., 2012), it remains to be seen whether the confluence of ER-Ca^2+^ gradients, CaMKII regulation of Arc localization and function, and the temporal dynamics of Arc could act as a mechanism for mGluR-LTD regulation and subsequent outputs. Furthermore, given that elevations of Arc are observed in several models of neuropsychiatric disease states along with dysregulation of LTD, determining the intracellular mechanisms that connect these phenomena is of great interest (Auerbach and Bear, 2010; Greer et al., 2010; Wu et al., 2011).

Here, we used the disrupted Arc ubiquitination mouse model (ArcKR) (Wall et al., 2018) to investigate these relationships. We found that hippocampal neurons isolated from ArcKR mice had an excess of ER-Ca^2+^ release upon chemical induction of mGluR-LTD in a subregion-dependent manner when compared to wildtype (WT) littermate control neurons. Overexpression of the ArcKR mutation in HEK293 cells increased Arc self-association compared to overexpressed WT Arc. Coexpression of ArcKR with CamKII in HEK293 cells also led to enhanced interactions with activated versions of CaMKII compared to wildtype (WT) Arc. Furthermore, Arc-CaMKII colocalization varied throughout the neuron, which was altered in ArcKR neurons. Collectively, our findings suggest a role for Arc ubiquitination in regulating ER Ca^2+^ dynamics, Arc self-association, and CaMKII signaling.

## 2 Methods

### 2.1 Animals

All animal care and use were carried out in accordance with the *National Institutes of Health Guidelines for the Use of Animals* using protocols approved by the Georgia State University Institutional Animal Care and Use Committee. Arc has been previously demonstrated to be ubiquitinated on K268 and K269 by the ubiquitin ligases RNF216 and UBE3A (Greer et al., 2010; Mabb et al., 2014). To mutate the ubiquitination sites of Arc, the conventional knock-in point mutation strategy was used where two point mutations were created within Exon 1 of the *Arc* gene resulting in the substitution of Lysine for Arginine at amino acid positions 268 and 269. The generation of this mouse did not disrupt Arc induction from its endogenous promoter but did selectively decrease the turnover of synthesized Arc protein due to its inability to be efficiently ubiquitinated on K268 and K269 (Wall et al., 2018). Homozygous Arc^WT/WT^ (WT) and Arc^KR/KR^ (KR) littermates were used for all experiments and were generated by crossing heterozygous Arc^WT/KR^ x Arc^WT/KR^ mice using the trio breeding method.

### 2.2 Cell culture

Primary hippocampal neuron cultures were generated from postnatal day 0-1 WT and ArcKR littermates as previously described (Wall et al., 2018). Neurons were grown on poly-d-lysine (0.1 mg/mL)-coated 6-well plates (9.6 cm^2^) at a density of 300,000 cells/well for Western blotting applications; on PDL-coated (1 mg/mL) coverslips (12 mm diameter) in 24-well plates (3.5 cm^2^) at a density of 75,000 cells/well for immunocytochemistry (ICC) applications; and on PDL-coated (1 mg/mL) coverslips (24 mm x 60 mm) in 60 mm culture dishes at a density of 75,000 cells/dish for live imaging experiments. Neurons were maintained in neuronal feeding media (Neurobasal media, ThermoFisher Scientific) containing 1% GlutaMAX (ThermoFisher Scientific), 2% B-27 (ThermoFisher Scientific), 4.8 μg/mL 5-Fluoro-2’-deoxyuridine (Sigma), and 0.2 ug/mL Gentamicin (Sigma) and fed by replacing half of the feeding media with fresh, pre-warmed neuronal feeding media every 3-4 days until used for experiments as described below.

HEK293 cells (ATCC) were seeded and maintained using standard procedures in Dulbecco’s modified Eagle’s medium (DMEM; Corning #10-017-CV) containing 10% FBS and 1% penicillinstreptomycin (ThermoFisher).

### 2.3 Highly Inclined Laminated Optical sheet (HILO) ER calcium imaging

Cultured primary hippocampal neurons were transfected with G-CatchER^+^ using Lipofectamine 2000 reagent (ThermoFisher Scientific) with a modified version of the manufacturer’s protocol as previously described (Reddish et al., 2021) at days in vitro (DIV) 11-12. Neurons were imaged at DIV 12-14.

To elicit and measure the magnitude of mGluR-mediated ER Ca^2+^ release, we used the type 1 mGluR agonist DHPG. G-CatchER^+^-transfected neurons were transferred to pre-warmed (37°C) artificial cerebrospinal fluid (ACSF; 124 mM NaCl, 3 mM KCl, 2 mM CaCl2, 2 mM MgCl2, 10 mM HEPES, 10 mM D-Glucose, pH 7.4). A G-CatchER^+^-positive neuron was identified and a signal was acquired at baseline. DHPG solution (100 μM (S)-3,5-DHPG (DHPG) (Tocris) in ACSF) was prewarmed to 37°C and then washed in manually using a syringe that was connected via tubing to the imaging chamber. After imaging, the DHPG was washed out with pre-warmed ACSF.

For HILO microscopy, a fiber-coupled 488 nm laser (Oxxius) was first collimated and then focused at the back focal plane of a 100X oil TIRF objective (N.A. 1.49, Nikon) using an achromatic lens with a focal length of 200 mm (Thorlabs) as previously described (Deng et al., 2021; Reddish et al., 2021). The position of the focused laser at the back focal plane of the objective was controlled by a 2D translational stage (Thorlabs). By laterally shifting the laser beam, the incident angle of laser beam at the interface of the coverslip and cell can be easily controlled. To achieve HILO imaging, the incident angle of the laser beam at the interface was adjusted to the sub-critical angle of total internal reflection where a thin optical light sheet was refracted into cells and excited the fluorophores within a few tens of micrometers from the coverslip surface. Laser power was kept consistent at 0.7 mW to minimize photobleaching. Images were acquired with a highly sensitive electron multiplying charge-coupled device (EMCCD) camera (Andor Ixon Ultra 888) at a rate of 2 frames/sec.

Live cell images were analyzed as previously described (Deng et al., 2021; Reddish et al., 2021) in ImageJ (NIH) by manually drawing a region of interest around clearly visible sub-regions of neurons (soma, primary branchpoints (PB)s, primary dendrites (PD)s, secondary branchpoints (SB)s, and secondary dendrites (SD)s) and fluorescence traces extracted. Raw pixel values were normalized to the average baseline fluorescence to generate a profile plot. Rigid motion correction was applied in cases where the neuronal subregion drifted partially or entirely out of the drawn regions of interest using the Rigid Registration ImageJ plugin. Videos were carefully inspected to verify cell health and data discarded in cases of clear cell morphological changes that indicated cell death (severe punctation, shriveling, or massive changes in fluorescence that were not associated with drug treatment).

Basal [Ca^2+^]_ER_ was calculated for WT and ArcKR neurons using a previously described method using pharmacological saturation (de Juan-Sanz et al., 2017; Reddish et al., 2021). Neurons were prepared and a signal was acquired as described above. Neurons were then treated with ACSF containing 50 μM ionomycin (ThermoFisher Scientific) in ACSF modified with 10 mM Ca^2+^. This saturates the G-CatchER^+^ signal, and using the known characteristics of the sensor, the following equation can be used to calculate the baseline ER Ca^2+^ concentration:

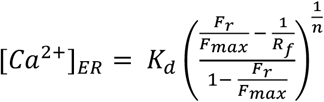

where K_d_ is the affinity coefficient of G-CatchER^+^, F_r_ is the average measured baseline fluorescence, R_f_ is the dynamic range of G-CatchER^+^, and n is the Hill coefficient.

### 2.4 Immunocytochemistry

Cultured primary hippocampal neurons on coverslips were fixed at DIV 12-14 with 4% paraformaldehyde and 4% sucrose in phosphate-buffered saline (PBS) at 37°C, washed with PBS, permeabilized with 0.1% Triton X-100 in PBS for 15 min at room temperature, and blocked with 10% normal goat serum (Gibco #16-210-072) for 1 hr at 37°C. Cells were then incubated with primary antibodies (rabbit anti-Arc (Synaptic Systems #156003; 1:500); rabbit anti-CaMKIIα/β (Abcam #ab52476; 1:250)) overnight at 4°C in 3% normal goat serum in PBS, washed with PBS, and incubated with secondary antibodies (donkey anti-mouse AlexaFluor 568 (Invitrogen #A10037; 1:1000); goat anti-rabbit AlexaFluor 488 (Life Tech #A11008; 1:1000)) for 1 hr at room temperature. After a final washing with PBS, coverslips were mounted onto slides with Fluorogel (GeneTex) or Vectashield Vibrance (Vector Laboratories) hardening mounting media with DAPI.

Neurons were imaged on a Zeiss LSM 700 confocal microscope using a 63X oil immersion objective lens (NA 1.4, Zeiss #420782-9900-000). Raw z-stack images (step size 0.38 μm) were analyzed using ImageJ (NIH) by manually drawing regions of interest around clearly visible sub-regions of neurons (soma, PBs, PDs, SBs, and SDs) and the integrated density was measured. All parameters related to acquisition were kept constant for all images within a given dataset. For colocalization analysis, the Coloc 2 plugin (bisection threshold regression) was used to calculate above-threshold pixel-pixel Pearson’s correlation coefficients between channels.

### 2.5 Western blotting

Cultured primary hippocampal neurons were harvested by briefly washing the plates in Dulbecco’s phosphate buffered saline (Gibco #14190144) and removed from the plate using a cell scraper (Corning #3010) and pelleted by centrifugation. Cell pellets were lysed in radioimmunoprecipitation assay (RIPA) buffer (150 mM NaCl, 50 mM Tris-HCl, 1% v/v Nonidet P-40, 0.05% sodium deoxycholate, 0.01% SDS) for 30 min on ice. For primary hippocampal neuron cultures, the entire cell lysate was used without assaying protein content due to low protein content and small volume. Following separation by SDS-PAGE, proteins were transferred to a nitrocellulose membrane (0.45 μm pore size, Bio-Rad). Membranes were blocked with Intercept TBS blocking buffer (LI-COR) overnight at 4°C and then incubated with primary antibodies (Arc, Synaptic Systems #156003, 1:1000; CaMKII, Abcam #ab52476, 1:1000; CaMKII p-Thr286 p-Thr287 Novus Biologicals #NB110-96896, 1:1,000; GAPDH, GeneTex #GTX627408, 1:1250) in a solution containing 1:1 blocking buffer:1% Tween-20 in tris-buffered saline (TBST) overnight at 4°C. After washing with TBST, membranes were incubated with secondary antibodies (IRDye Goat anti-Mouse 680RD, LI-COR #926-68070, 1:15,000, IRDye Goat anti-Rabbit 800CW, LI-COR #926-32211, 1:15,000) in 1:1 blocking buffer:TBST for 1 hr at room temperature. Membranes were imaged on a LI-COR Odyssey CLx scanner (low scan quality, 163 μm scan resolution, auto channel intensities) and resulting images were analyzed using the gel analysis tool in ImageJ (NIH). For images where high background was present, background subtraction was performed on the whole channel using the Subtract Background tool in ImageJ with a rolling ball radius of 20-50 pixels.

### 2.6 Co-immunoprecipitation

HEK293 cells were seeded on 6-well plates and transfected with pRK5-myc-Arc, pRK5-myc-ArcKR, pEGFP-Arc, pEGFP-ArcKR, pCAG-mEGFP-CaMKIIα, and/or pCAG-mEGFP-CaMKIIβ, pCAG-mEGFP-CaMKIIα T286A, pCAG-mEGFP-CaMKIIα T286D, pCAG-mEGFP-CaMKIIβ T287A, pCAG-mEGFP-CaMKIIβ T287D, and/or mVenus-Calnexin (Addgene #56324) (CaMKII plasmids were a kind gift from Dr. Ulli Bayer at the University of Colorado Medicine (Bayer et al., 2006b)) using Lipofectamine 2000 or Lipofectamine 3000 reagent (ThermoFisher Scientific) following the manufacturer procedure. After allowing constructs to express overnight, cells were harvested and lysed in immunoprecipitation (IP) buffer (20 mM Tris-HCl, 3 mM EDTA, 3 mM EGTA, 150 mM NaCl, 1% Triton X-100, 1 mM DTT, pH 7.4) containing protease and phosphatase inhibitors (0.1 mM phenylmethylsulfonyl fluoride, 1 μM leupeptin, 0.15 μM aprotinin, and 1:2000 Halt phosphatase inhibitor cocktail (ThermoFisher Scientific #78420)) for 30 min on ice. After centrifugation (14,000 rpm, 25 min, 4°C), lysate protein concentration was determined using a Pierce 660 assay (ThermoFisher). 10 μg of protein was used for subsequent steps, with 1 μg of protein (10%) reserved for input control. Lysates were incubated with 0.5 μg/mL monoclonal mouse anti-myc antibody (Santa Cruz Biotech #sc-40) for 1 hr at 4°C. 25 μL of pre-equilibrated GammaBind Sepharose beads (Cytiva) were then added and samples tumbled overnight at 4°C. Samples were washed with IP buffer and proteins were eluted from beads using SDS sample buffer (Li-Cor #928-40004) with heating at 45°C for 5 min. Subsequently, beads were pelleted by centrifugation (15,000 rpm for 2 min) and the supernatant containing protein was separated using SDS-PAGE, transferred to a nitrocellulose membrane, and immunoblotting proceeded as described above using anti-GFP (Fisher #NB600308, 1:1000) and polyclonal goat anti-myc (Bethyl/ThermoFisher #A190-104A, 1:1000) primary antibodies. For quantification, GFP-tagged co-immunoprecipitated protein band intensities were normalized to the immunoprecipitated myc-tagged protein, which served as an important internal control for IP efficiency.

### 2.7 Arc protein purification and test of oligomerization

Purification of His-tagged ArcWT and ArcKR was carried out by a third-party company (GenScript Biotech, New Jersey, USA) using the HD transient expression of recombinant protein – gold system. Briefly, the sequences for ArcWT and ArcKR were synthesized and cloned into the pcDNA3.4 target vector using the EcoRI and HindIII restriction sites. Plasmids were then transfected into the HD 293F cell line for expression. Proteins were purified using HisTrap FF Crude (Cytiva) purification columns and eluted in PBS, pH 7.2. Purity was too low to be detected by SDS-PAGE under nonreducing conditions. After receipt, protein was further concentrated using Amicon Ultra-0.5 mL centrifugal filters (Millipore Sigma) prior to use. Proteins were brought up in the same concentration and used for subsequent SDS-PAGE analysis under reducing (SDS sample buffer containing 5% 2-mercaptoethanol (2-ME)) and nonreducing conditions (SDS sample buffer without 2-ME) after heating for 5 min at 95°C.

### 2.8 Statistical analysis

Data are presented as mean +/- SEM unless otherwise indicated. Statistical tests were performed using GraphPad Prism software. Statistical significance was defined as p < 0.05. For testing one independent variable when experimental groups had equal variance as determined using and F test, a t-test was used; otherwise, a t-test was used with Welch’s correction. For comparing two independent variables, a two-way ANOVA was used. To determine differences between groups following a two-way ANOVA, either Sidak’s test for multiple comparisons or Tukey’s test was used depending on the relevant comparisons. The precise statistical test used for each experiment, including sample sizes, factors, and post-hoc comparisons, can be found in the Results section describing the experiment.

## 3 Results

### 3.1 Excess ER calcium release in ArcKR primary hippocampal neurons

In our previous studies, CA1 field recordings from acute hippocampal slices showed that ArcKR neurons displayed enhanced mGluR-LTD in response to application of S-3,5-dihydroxyphenylglycine (DHPG), including a reduced threshold for reaching LTD and a higher magnitude of LTD (Wall et al., 2018). Because mGluR-LTD is a G_q_ dependent process requiring release of intracellular Ca^2+^ stores from the ER via ER membrane-bound IP_3_Rs, we first explored the possibility that DHPG elicits altered ER Ca^2+^ release kinetics in ArcKR neurons.

ArcWT (WT) and ArcKR (KR) primary hippocampal neuron cultures were transfected with the ER-localized genetically-encoded calcium indicator G-CatchER^+^ (Reddish et al., 2021) and imaged on a highly inclined laminated optical sheet (HILO) microscope during the addition of 100 μM S-3,5-DHPG. G-CatchER^+^ is specifically targeted and expressed in the ER lumen; thus, a decrease in G-CatchER^+^ fluorescence reflects Ca^2+^ being released from the ER (Reddish et al., 2021). As the subcellular distribution of Ca^2+^ signaling-related proteins, ER complexity, intracellular [Ca^2+^], and ER Ca^2+^ release can vary throughout the neuron (Sharp et al., 1993; Terasaki et al., 1994; Spacek and Harris, 1997; Blaustein and Golovina, 2001; Cui-Wang et al., 2012; Krijnse-Locker et al., 2017), we examined various neuronal subregions separately: soma, primary branchpoints (PBs), primary dendrites (PDs), secondary branchpoints (SBs), and secondary dendrites (SDs) (Deng et al., 2021; Reddish et al., 2021) (Figures 1A-1C). G-CatchER^+^ fluorescence traces showed a striking depletion of ER Ca^2+^ stores in KR neurons after DHPG application (Figure 1). ER Ca^2+^ release as quantified by percent decrease from baseline average was significantly enhanced in KR neurons at the soma (Figures 1D,E; two-way ANOVA, Genotype: F(1, 32) = 20.74, p < 0.0001, Time: F(1, 32) = 10.63, p = 0.0026, Interaction: F(1, 32) = 18.49, p = 0.0001; Sidak’s post-hoc test, WT vs KR at End p < 0.0001), PB (Fig. 1F,G; two-way ANOVA, Genotype: F(1, 32) = 9.120, p = 0.0049, Time: F(1, 32) = 39.59, p < 0.0001, Interaction: F(1, 32) = 6.651, p = 0.0147; Sidak’s post-hoc test, WT vs KR at End p = 0.0008), PD (Figures 1H,I; two-way ANOVA, Genotype: F(1, 32) = 7.966, p = 0.0081, Time: F(1, 32) = 24.04, p < 0.0001, Interaction: F(1, 32) = 7.966, p = 0.0063; Sidak’s post-hoc test, WT vs KR at End p = 0.0006), and SD (Figures 1L,M; two-way ANOVA, Genotype: F(1, 32) = 32.51, p < 0.0001, Time: F(1, 32) = 36.77, p < 0.0001, Interaction: F(1, 32) = 48.36, p < 0.0001; Sidak’s post-hoc test, WT vs KR at End p < 0.0001). However, KR SB showed no difference in release compared to WT neurons (Figures 1J,K; two-way ANOVA, Genotype: F(1, 32) = 0.3066, p = 0.5836, Time: F(1, 32) = 8.865, p = 0.0055, Interaction: F(1, 32) = 0.6407, p = 0.4294), which related to our previous findings that Ca^2+^ release at secondary branchpoints is reduced (Deng et al., 2021; Reddish et al., 2021).

**Figure 1.**
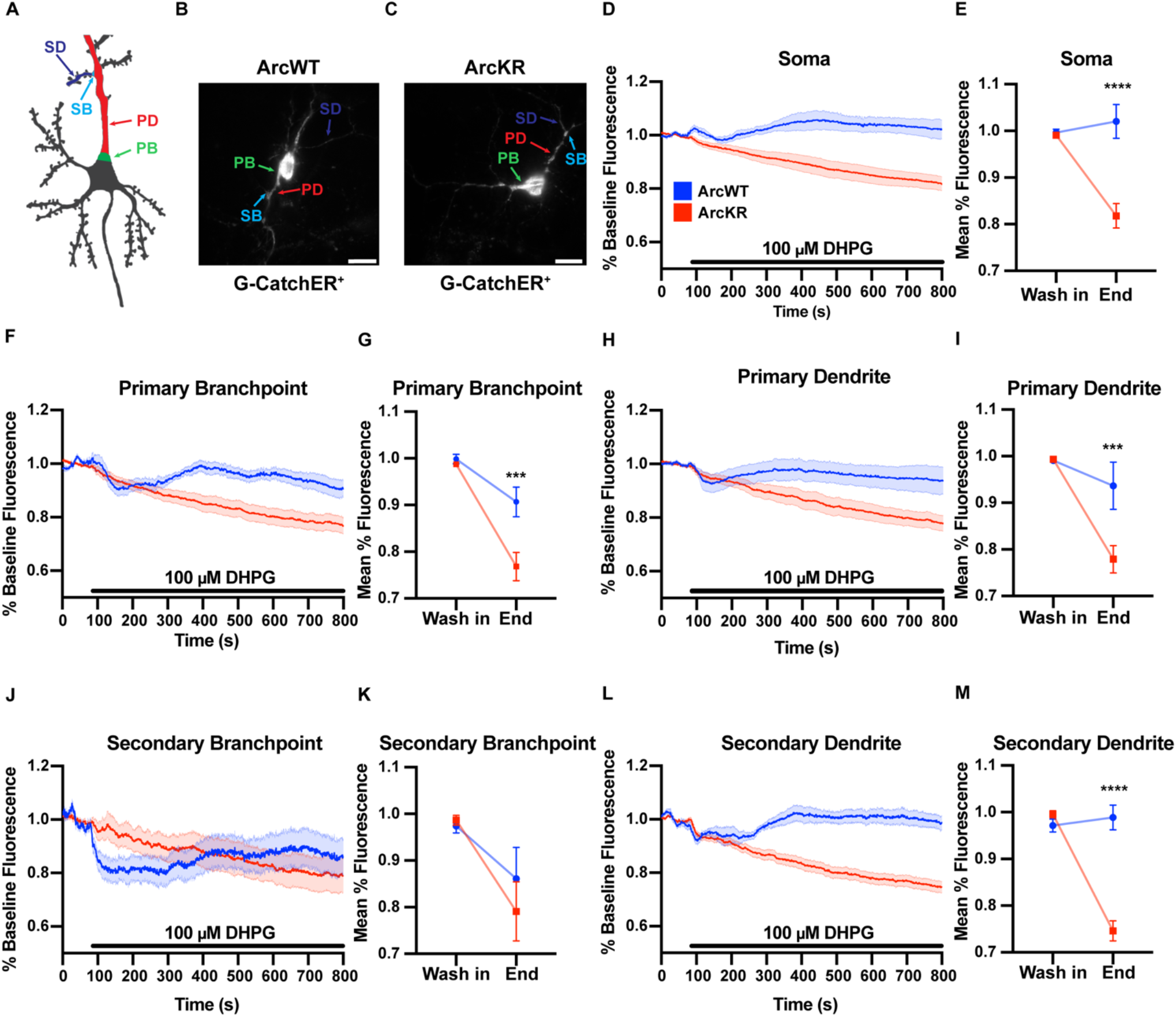
DHPG elicits excess ER Ca^2+^ release with disrupted dynamics in ArcKR hippocampal neurons. **(A)** Schematic of regions of interest referred to in this figure. **(B)** Representative image from a G-CatchER^+^-transfected ArcWT neuron (**C**) Representative image from a G-CatchER^+^-transfected ArcKR neuron. **(D, F, H, J, L)** Normalized G-CatchER^+^ signal for WT (blue, N = 6 neurons) and ArcKR (red, N = 12 neurons) primary hippocampal neurons in the region of the neuron indicated with application 100 μM DHPG (black bar). Mean percent change from baseline (0-75 s) +/- SEM. Soma n = 6 WT, 12 KR; primary branchpoint n = 13 WT, 25 KR; primary dendrite n = 6 WT, 12 KR; secondary branchpoint n = 16 WT, 14 KR; secondary dendrites n = 14 WT, 24 KR. **(E, G, I, K, M)** Mean G-CatchER^+^ fluorescence decrease from mean baseline fluorescence (0-75 s) at wash-in of DHPG and at 800 s ArcWT (blue) and ArcKR (red) primary hippocampal neurons. Mean +/- SEM. Two-way ANOVA with Šídák’s multiple comparison test; * p < 0.05, ** p < 0.01, *** p < 0.001, **** p < 0.0001, ns p > 0.05.

To evaluate if the enhanced Ca^2+^ release in ArcKR neurons was simply due to baseline increases in ER Ca^2+^, we measured baseline [Ca^2+^]_ER_ during the application of the ionophore ionomycin (50 μM) in a buffer containing excess Ca^2+^ (10 mM) to saturate the G-CatchER^+^ signal and [Ca^2+^] was calculated as previously described (de Juan-Sanz et al., 2017) (Figure 2). However, we found no genotype differences in ER Ca^2+^ in any neuronal region examined (Figure 2F; two-way ANOVA, Genotype: F(1, 41) = 1.133, p = 0.2935, Region: F(4, 41) = 7.998, p < 0.0001, Interaction: F(4, 41) = 0.2939, p = 0.8803), indicating that the phenotypes observed in KR neurons were not influenced by baseline [Ca^2+^]_ER_ prior to DHPG application.

**Figure 2.**
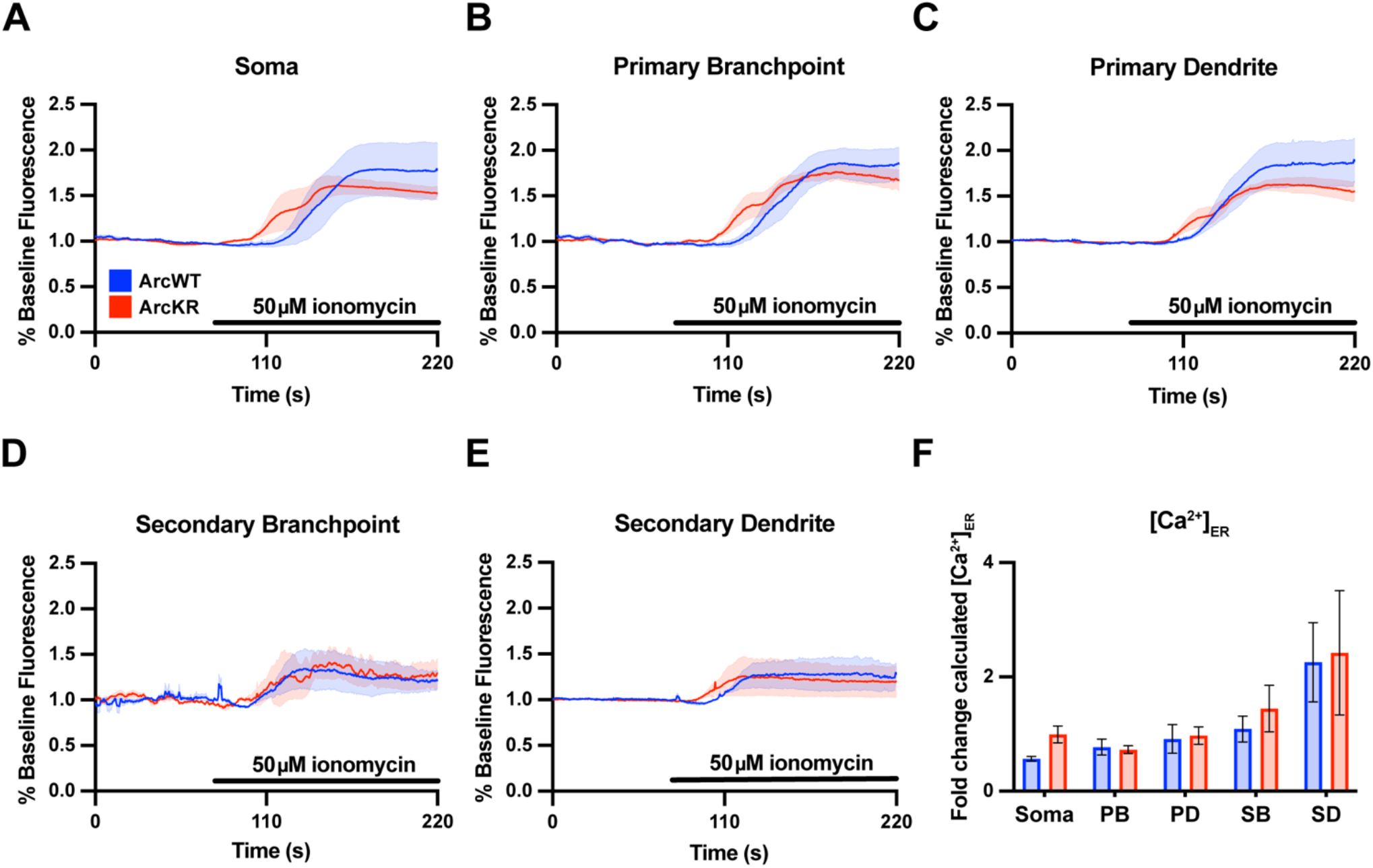
Baseline ER [Ca^2+^] is not different in ArcKR primary hippocampal neurons. **(A-E)** Mean G-CatchER^+^ fluorescence change from baseline (0-75 s) at the indicated regions during application of 50 μM ionomycin + 10 mM Ca^2+^ to ArcWT (blue, N = 3) and ArcKR (red, N = 4) primary hippocampal neurons. Mean +/- SEM. Soma n = 3 WT, 4 KR; primary branchpoint n = 8 WT, 8 KR; primary dendrite n = 8 WT, 8 KR; secondary branchpoint n = 4 WT, 3 KR; secondary dendrites n = 3 WT, 2 KR. **(F)** Calculated [Ca^2+^]_ER_ for ArcWT and ArcKR neurons. Mean +/- SEM. Two-way ANOVA with Šídák’s multiple comparison test, no significant genotype effects. Significant simple effect of region (F (4, 41), p < 0.0001).

### 3.2 ArcKR displays altered interaction with CaMKII that depends on CaMKII isoform- and phosphorylation status

Previous work has shown that Arc interacts directly with CaMKII, and that this interaction depends on the isoform and phosphorylation status of CaMKII (Okuno et al., 2012; Zhang et al., 2015, 2019). We transfected HEK293 cells with either myc-ArcWT or myc-ArcKR along with GFP-CaMKIIa or GFP-CaMKIIb. We also used CaMKII plasmids with point mutations at Thr286 (for CaMKIIa) or Thr287 (CaMKIIb) that mimic the active, autophosphorylated state (T286D/T287D) or mimic the inactive unphosphorylated state (T286A/T287A) (Bayer et al., 2006a; O’Leary et al., 2006; Coultrap et al., 2012). After immunoprecipitation (IP) of Arc, protein complexes were eluted and immunoblotting proceeded for myc- and GFP-tagged proteins (Figures 3A,C,E,G,I).

**Figure 3.**
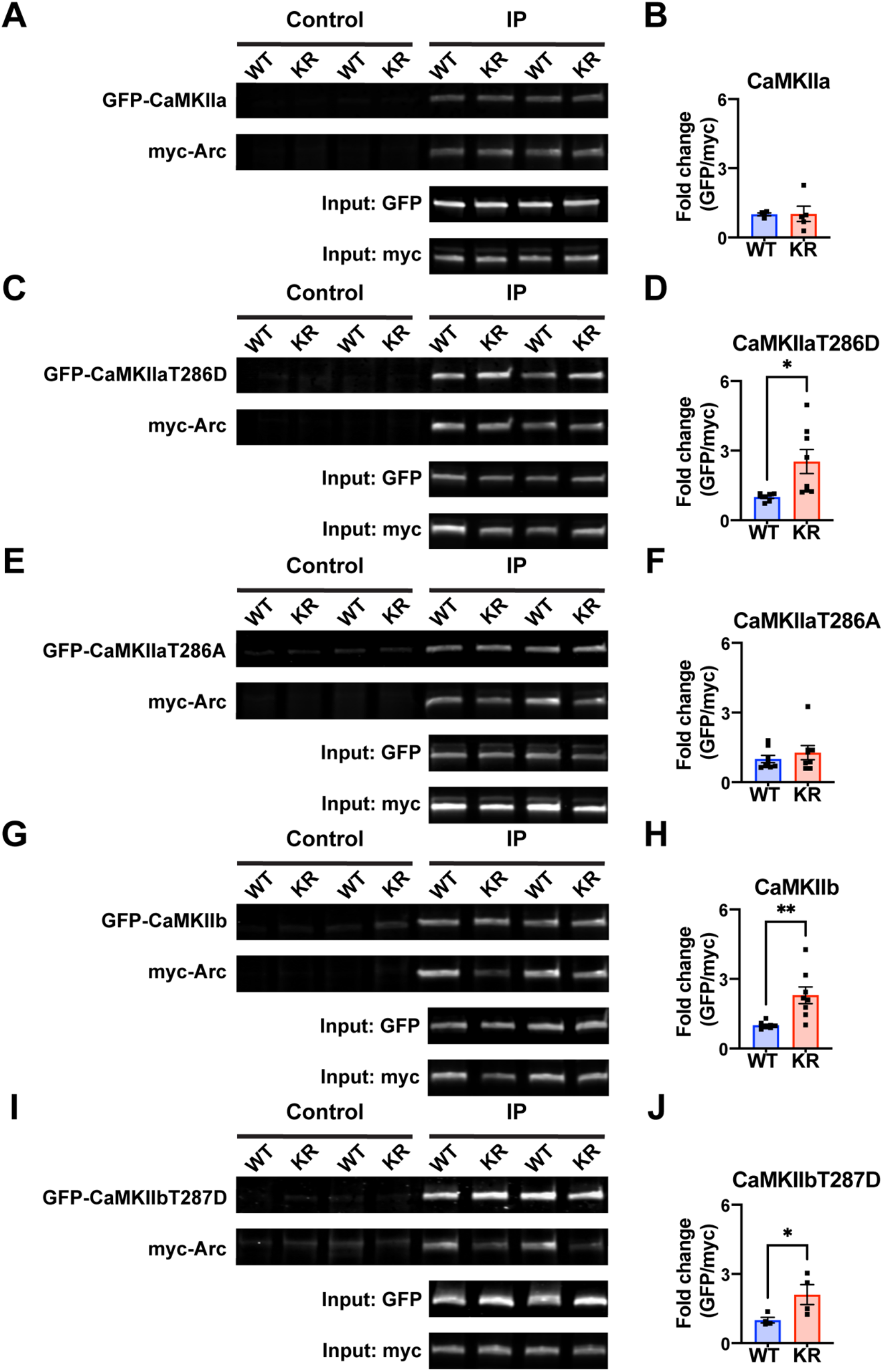
ArcKR protein differentially interacts with CaMKII in an isoform- and phosphorylation status-dependent manner. **(A, C, E, G, I)** Representative Western blots after pulldown of myc-Arc and immunoblotted for the indicated tags. **(B)** Quantification of GFP-CaMKIIa after IP myc-ArcWT (blue) or myc-ArcKR (red). **(D)** Quantification of GFP-CaMKIIaT286D after IP of myc-ArcWT or myc-ArcKR. Unpaired t-test with Welch’s correction for unequal variances, t = 2.957, df = 7.210, * p = 0.0205. **(E)** Quantification of GFP-CaMKIIaT286A after IP of myc-ArcWT or myc-ArcKR. Unpaired t-test with Welch’s correction for unequal variances, t = 0.8050, df = 14, p = 0.4343. **(H)** Quantification of GFP-CaMKIIb after IP of myc-ArcWT or myc-ArcKR. Unpaired t-test with Welch’s correction for unequal variances, t = 3.564, df = 7.273, ** p = 0.0086. **(J)** Quantification of GFP-CaMKIIbT287D after IP of myc-ArcWT or myc-ArcKR. Unpaired t-test, t = 2.454, df = 6, * p = 0.0495. Data are represented as mean fold change relative to WT +/- SEM.

While CaMKIIa-GFP interaction with ArcKR was not statistically different from WT (Figure 3B; unpaired t-test with Welch’s correction, t = 0.0607, df = 4.289, p = 0.9543), ArcKR interaction was significantly higher with CaMKIIaT286D (Figure 3D; unpaired t-test with Welch’s correction, t = 2.957, df = 7.210, p = 0.0205). No difference was detected with CaMKIIaT286A (Figure 3F; unpaired t-test, t = 0.8050, df = 14, p = 0.4343). Interestingly, ArcKR interaction with CamKIIb was significantly higher than WT for both WT CaMKIIb as well as CaMKIIbT287D (Figures 3H,J; CaMKIIb unpaired t-test with Welch’s correction, t = 3.564, df = 7.273, p = 0.0086, CaMKIIbT287D unpaired t-test, t = 2.454, df = 6, p = 0.0495). These results indicate that ArcKR interacts more with CaMKIIb, and that autophosphorylation status of CaMKIIa and CaMKIIb could determine ArcKR interaction.

### 3.3 Overexpressed and purified ArcKR protein exhibits increased self-association

Arc oligomerization has been described to be regulated by Ser-260 phosphorylation by CaMKII (Zhang et al., 2019). Because the mutated lysines of ArcKR (K268/269) are in close proximity to Ser-260 (Mabb et al., 2014; Wall et al., 2018), we sought to determine whether Arc oligomerization potential is altered. We co-transfected HEK293 cells with N-terminal myc-tagged Arc (myc-ArcWT) or ArcKR (myc-ArcKR) and N-terminal GFP-tagged Arc (GFP-ArcWT) and GFP-ArcKR and performed a co-immunoprecipitation assay to compare the amount of self-association.

Immunoblotting following immunoprecipitation of myc-Arc and myc-ArcKR showed that ArcKR self-associates more than ArcWT (Figures 4A,B; unpaired t-test with Welch’s correction, t = 3.336, df = 8.199, p = 0.0099). Greater self-pulldown does not indicate oligomerization *per se*, as this method does not differentiate between direct and indirect interactions. Therefore, we next purified ArcWT and ArcKR from HEK 293 cells and performed SDS-PAGE on the purified His-ArcWT and His-ArcKR proteins under reducing and non-reducing conditions (Figures 4C). Under non-reducing conditions, we observed high molecular weight reactive Arc species in both ArcWT and ArcKR. However, there was a prominent band between 100 and 150 kDa in ArcWT but not ArcKR indicating that a fraction of the ArcWT can exist as a dimer (blue asterisk). Under reducing conditions, both ArcWT and ArcKR collapsed into their monomeric forms with prominent bands between 50 and 75 kDa (white asterisk). Cumulatively, these findings suggest that ArcKR is more predisposed to forming higher molecular weight oligomers compared to ArcWT.

**Figure 4.**
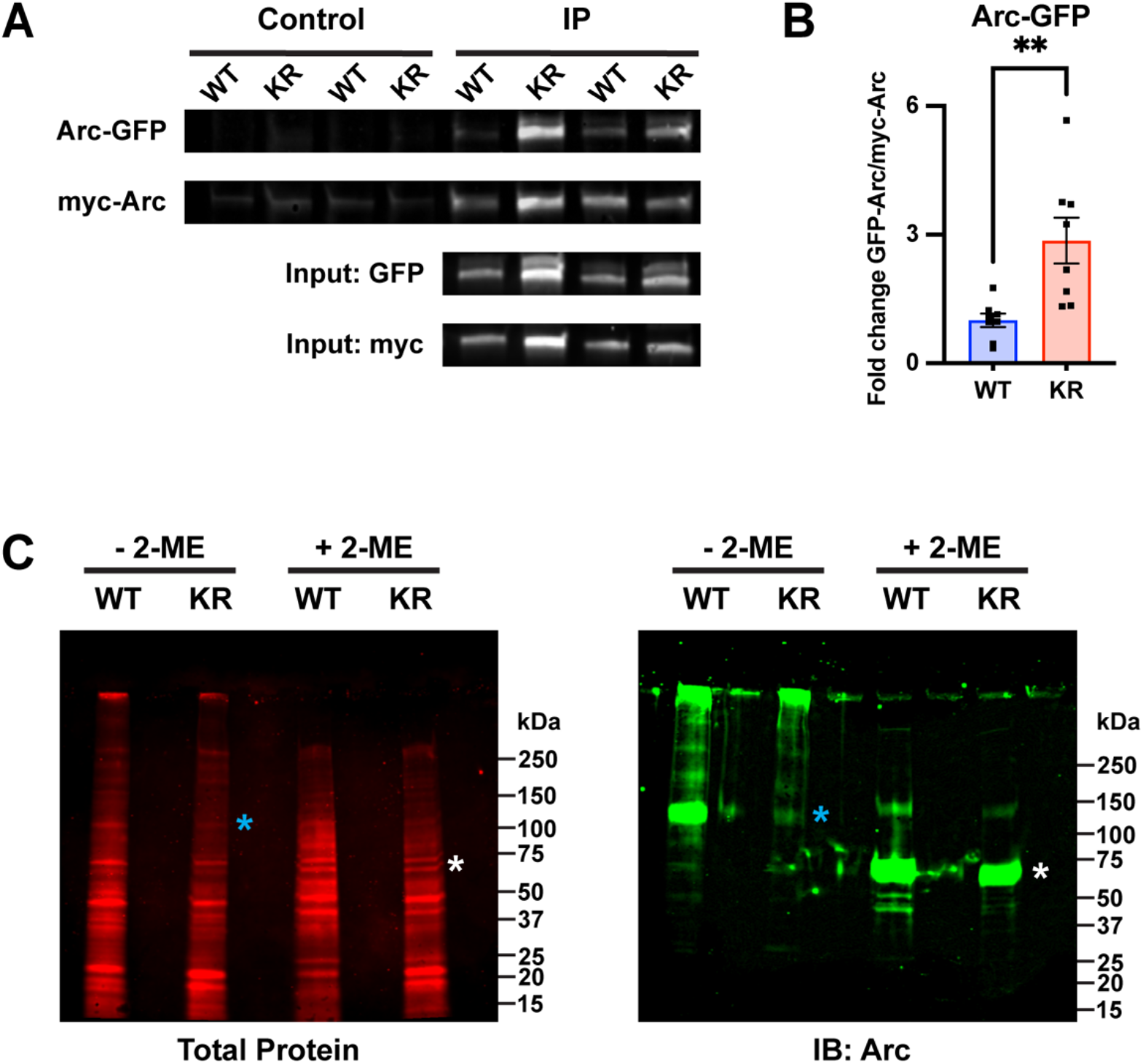
ArcKR self-association is increased relative to ArcWT. **(A)** Representative Western blots after pulldown of myc-ArcWT or myc-ArcKR with an anti-myc antibody and immunoblotted with anti-GFP to detect GFP-ArcWT or GFP-ArcKR. **(B)** Quantification of GFP-ArcWT or GFP-ArcKR after pulldown with myc-Arc or myc-ArcKR. t-test with Welch’s correction for unequal variances, t = 3.336, df = 8.199, ** p = 0.0099. Mean fold change relative to WT +/- SEM. **(C)** His-ArcWT and His-ArcKR under non-reducing (−2-ME) and reducing conditions (+ 2-ME). *Left*, total protein stain for Arc. *Right*, Immunoblot using an anti-Arc polyclonal antibody (blue * represents predicted Arc dimer, white * represents predicted Arc monomer).

### 3.4 ArcKR neurons display regional alterations in Arc-CaMKII colocalization

Arc and CaMKII localization and colocalization within subregions of the neuron are critical for several signaling pathways that support learning and memory (Davies et al., 2007; Marsden et al., 2010), including localizing Arc to inactive synapses (Okuno et al., 2012). Given that our previous studies on Arc-CaMKII interactions were conducted in HEK293 cells, we sought to establish relationships of endogenously expressed Arc and CaMKII in primary hippocampal neurons. To discern potential changes in CaMKII localization and colocalization with Arc, we immunostained ArcWT and ArcKR primary hippocampal neuron cultures for CaMKII and Arc and imaged their distributions using confocal microscopy (Figures 5-6).

**Figure 5.**
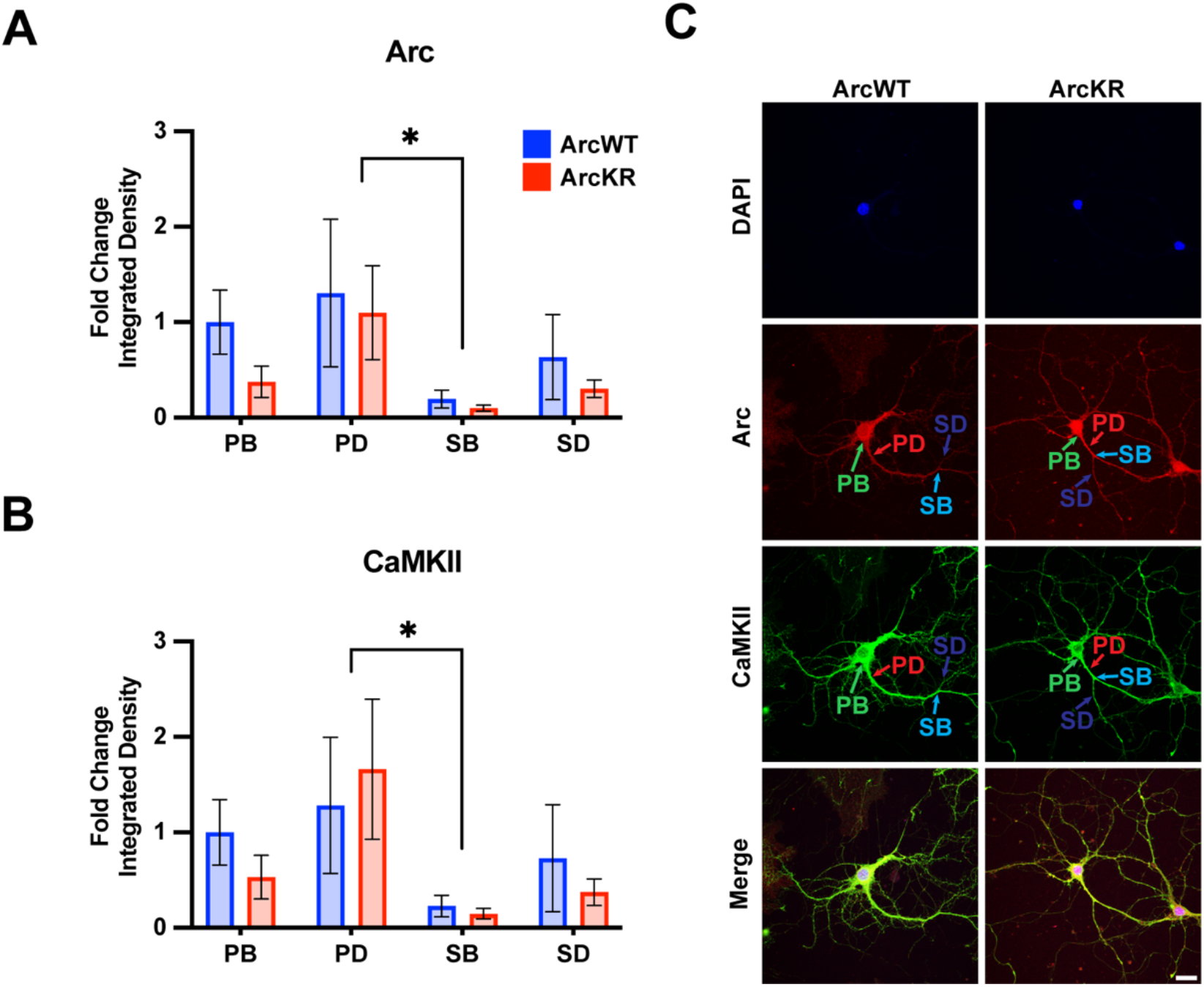
Arc and CaMKII are decreased at secondary branchpoints in ArcWT and ArcKR primary hippocampal neurons. **(A)** Fold change of Arc integrated densities for the indicated regions in ArcWT (blue, N = 7) and ArcKR (red, N = 5) primary hippocampal neurons relative to ArcWT PB. Primary branchpoint n = 12 WT, 11 ArcKR; primary dendrite n = 9 WT, 10 ArcKR; secondary branchpoint n = 10 WT, 11 ArcKR; secondary dendrites n = 10 WR, 11 ArcKR. Tukey’s post-hoc test, PD vs SB: *p = 0.0270. **(B)** CaMKII integrated densities for indicated regions in (A). Tukey’s post-hoc test PD vs SB: *p = 0.0187. **(C)** Representative confocal images from ArcWT and ArcKR primary hippocampal neurons. Scale bar = 20 μm. Data represented as mean fold change relative to WT PB +/- SEM.

**Figure 6.**
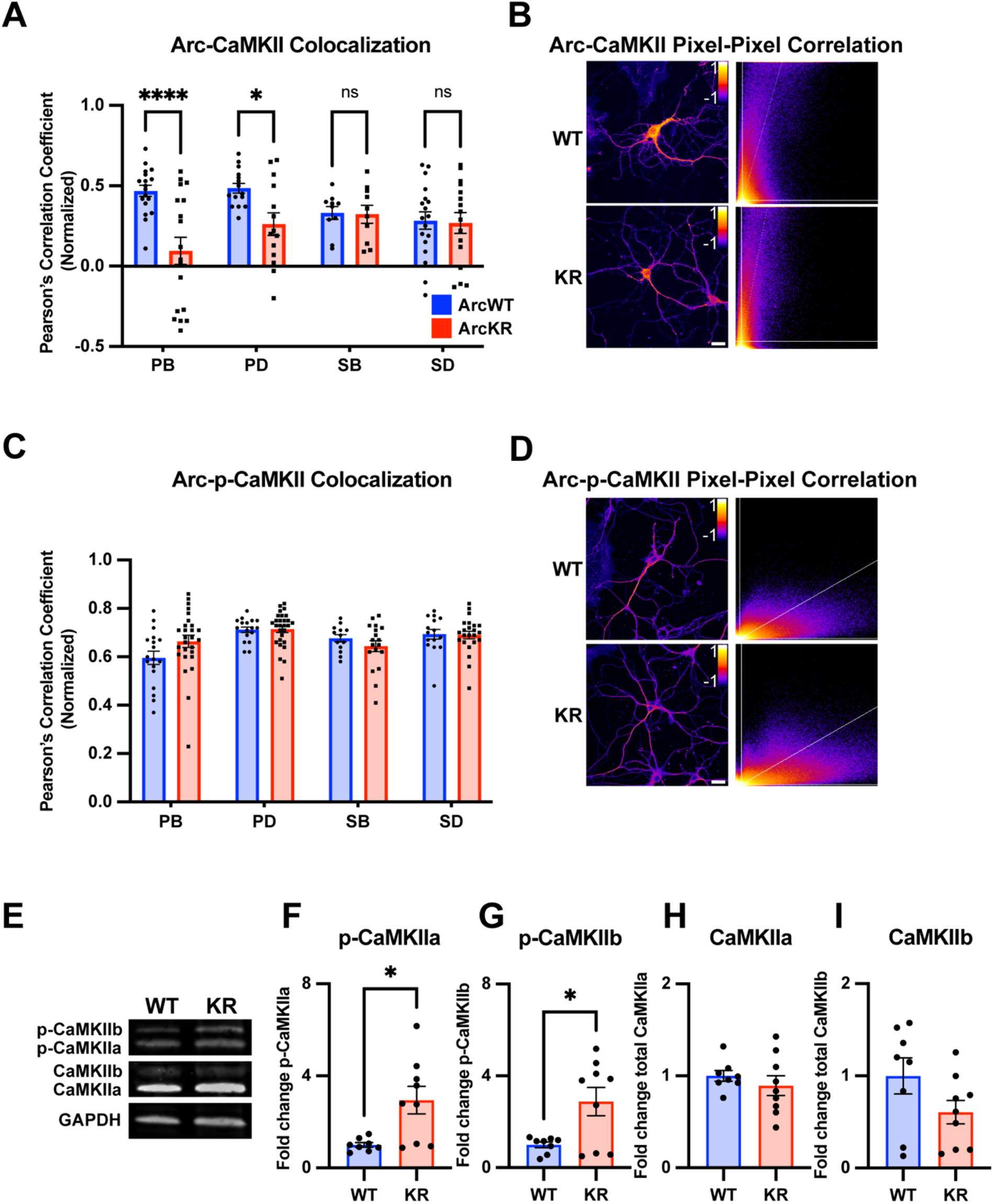
Global and regional alterations in Arc-CaMKII colocalization in ArcKR hippocampal neurons. **(A)** Pearson correlation coefficient between CaMKII and Arc in ArcWT (blue, N = 7) and ArcKR (red, N = 5) primary hippocampal neurons. Primary branchpoint n = 18 WT, 18 KR; primary dendrite n = 15 WT, 14 KR; secondary branchpoint n = 10 WT, 10 KR; secondary dendrites n = 18 WT, 16 KR. Twoway ANOVA with Šídák’s multiple comparison test, effect of genotype (F(1, 111) = 12.54, p < 0.0006, interaction F(3,111) = 4.475, p = 0.0053). * p < 0.05, **** p < 0.0001, ns p > 0.05. **(B)** Representative pixel-pixel correlation heatmap between Arc and CaMKII for an ArcWT and ArcKR neuron (left) and corresponding pixel-pixel intensity plots with trendline (right). Scale bar = 20 μm. **(C)** Pearson correlation coefficient between p-CaMKII and Arc in ArcWT (blue, N = 7) and ArcKR (red, N = 9) primary hippocampal neurons. Primary branchpoint n = 18 WT, 27 KR; primary dendrite n = 17 WT, 28 KR; secondary branchpoint n = 13 WT, 18 KR; secondary dendrites n = 15 WT, 23 KR. Two-way ANOVA with Šídák’s multiple comparison test. **(D)** Representative pixel-pixel correlation heatmap between Arc and p-CaMKII for a WT and KR neuron (left) and corresponding pixel-pixel plots with trendline (right). Scale bar = 20 μm. **(E)** Representative Western blots for (F-I). **(F)** Quantification of Western blot for p-CaMKIIα normalized to total CaMKIIα. Unpaired t-test (t = 3.015, df = 15, ** p = 0.0087). **(G)** Quantification of Western blot for p-CaMKIIβ normalized to total CaMKIIβ. Unpaired t-test (t = 2.821, df = 15, * p = 0.0129). **(H)** Quantification of Western blot for total CaMKIIα. Unpaired t-test (t = 0.8324, df = 15, p = 0.4183). **(I)** Quantification of Western blot for total CaMKIIβ. Unpaired t-test (t = 1.712, df = 15, p = 0.1075). For (A) and (C), data represented as mean fold change relative to WT PB +/- SEM, for F-I data are represented as mean fold change relative to WT +/- SEM.

Two-way ANOVA found no significant genotype difference in Arc protein levels (Figure 5A; two-way ANOVA, Genotype: F(1, 76) = 1.535, p = 0.2192, Region: F(3, 76) = 2.890, p = 0.0409, Interaction: F(3, 76) = 0.2099, p = 0.8892); however, there was a significant effect of region, with secondary branchpoints displaying lower levels than primary dendrites (Tukey’s post-hoc test, PD vs SB: p = 0.0270). Similarly, CaMKII levels were lower in the secondary branchpoints compared to primary dendrites (Figure 5B; two-way ANOVA, Genotype: F(1, 76) = 0.3949, p = 0.7570, Region: F(3, 76) = 3.160, p = 0.0294, Interaction: F(3, 76) = 0.1961, p = 0.6592, Tukey’s post-hoc test PD vs SB: p = 0.0187).

We next quantified the colocalization of Arc with total CaMKII from the same samples used in Figure 5. Pixel-pixel Pearson correlation values were lower in KR neurons compared to WT in PBs and PDs but not SBs or SDs (Figure 6A-B; two-way ANOVA, Genotype: F(1, 111) = 12.54, p = 0.0006, Region: F(3, 111) = 1.166, p = 0.3261, Interaction: F(3, 111) = 4.475, p = 0.0053; Sidak’s post-hoc test WT vs KR PB p < 0.0001, PD p = 0.0425, SB p > 0.9999, SD p = 0.9995). Given the increase in ArcKR interaction with both CamKIIa and CamKIIb constitutive active mutants, we stained ArcWT and ArcKR hippocampal neuron cultures for p-CaMKII and calculated the Arc-p-CaMKII colocalization. However, there were no significant genotype differences (Figure 6C-D; two-way ANOVA Genotype: F(1, 151) = 0.3609, p = 0.5489, Region: F (3, 151) = 6.628, p = 0.0003, Interaction: F (3, 151) = 2.056, p = 0.1085; Sidak’s post-hoc test, PB vs PD p = 0.0003, PB vs SD p = 0.0181).

Because of constraints in resolving isoform-specific changes in CaMKII levels with immunostaining, we used Western blotting to determine if CaMKIIa and CaMKIIb along with their respective levels of phosphorylation were altered in ArcKR hippocampal neuron cultures (Figure 6E). We observed that the proportion of the phosphorylated forms of CaMKIIa (p-CaMKIIa) and CaMKIIb (p-CaMKIIb) were greater in ArcKR neurons (Figure 6F-G; p-CaMKIIa unpaired t-test, t = 3.015, df = 15, p = 0.0087; p-CaMKIIb unpaired t-test, t = 2.821, df = 15, p = 0.0129) but there was no significant difference in the total expression of either isoform (Figures 6H-I; CaMKIIa unpaired t-test, t = 0.8324, df = 15, p = 0.4183; CaMKIIb unpaired t-test, t = 1.712, df = 15, p = 0.1075). Taken together, these results suggest that while ArcKR protein and total CaMKII levels are unchanged in ArcKR neurons, ArcKR is biased toward interacting with p-CaMKII that may occur in a region-dependent manner.

### 3.5 ArcKR has greater interaction with the ER-bound protein calnexin

As we are not aware of any literature implicating Arc levels or posttranslational modifications in regulating ER-mediated Ca^2+^ release, we performed co-immunoprecipitation assays with proteins known to be important for CaMKII or ER function. First, we considered the calcium-sensitive protein calmodulin which is required for CaMKII autophosphorylation (Lisman et al., 2002). However, we were unable to detect an interaction of GFP-tagged calmodulin with ArcWT or ArcKR (Figure 7A).

**Figure 7.**
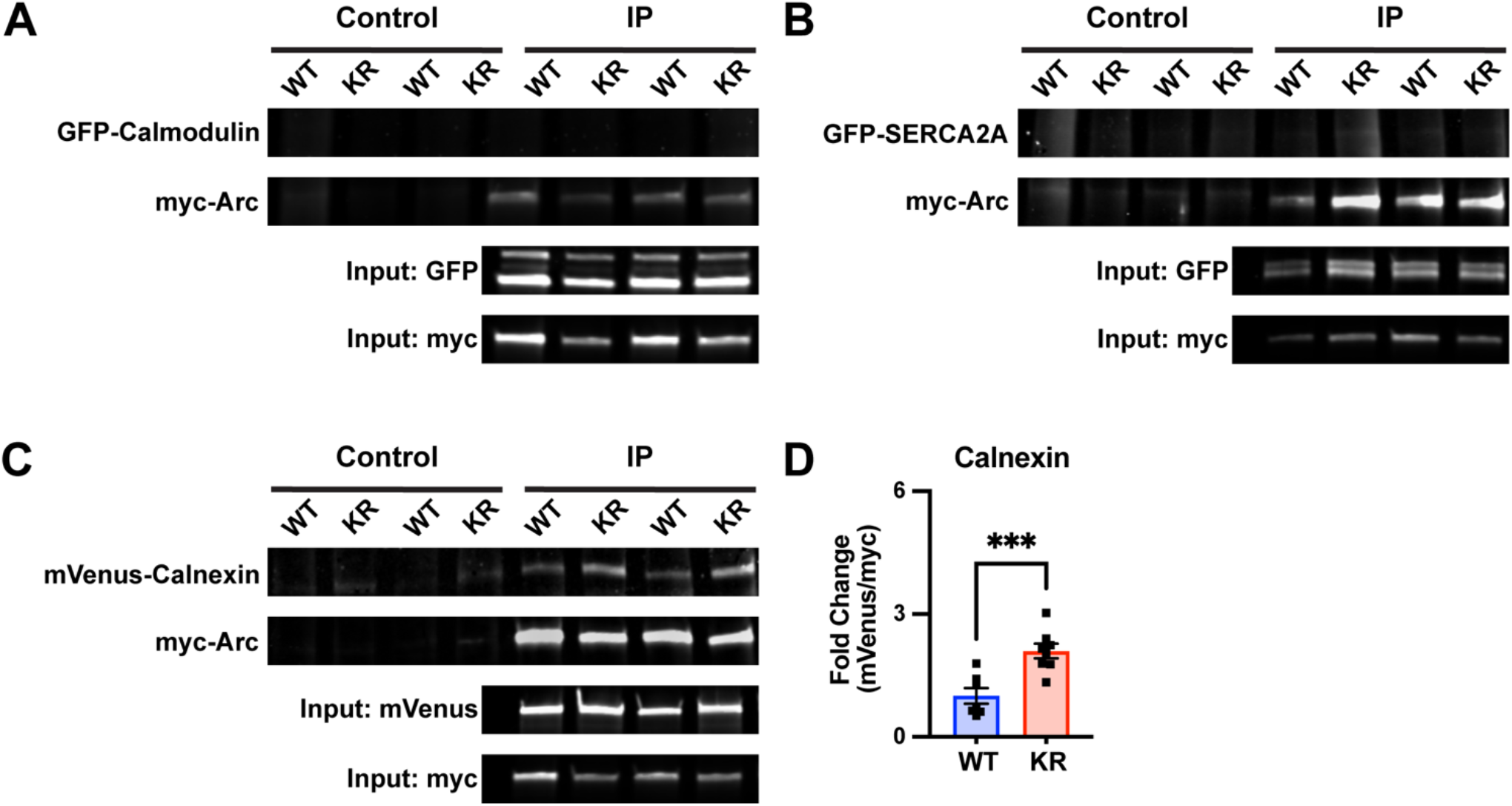
Enhanced interaction of ArcKR with the ER membrane protein Calnexin. **(A-C)** Representative blots after pulldown of myc-ArcWT or myc-ArcKR and immunoblotted for the indicated tagged proteins. **(D)** Increased co-immunoprecipitation of Calnexin with ArcKR (Unpaired t-test, p = 0.0009). Mean fold change relative to WT +/- SEM.

Arc has also been shown to directly interact with the cytoplasmic tail of the ER membrane-bound protein calnexin, although the functional consequence of this interaction is unknown (Myrum et al., 2017). Intriguingly, calnexin regulates the activity of the sarco/endoplasmic reticulum Ca^2+^-ATPase (SERCA), another ER-bound protein that sequesters Ca^2+^ in the lumen of the ER (Britzolaki et al., 2020). While SERCA2A did not co-immunoprecipitate with either ArcWT or ArcKR (Figure 7B), coimmunoprecipitation in HEK293 cells reaffirmed interaction of Arc with calnexin and, notably, revealed that ArcKR protein has an enhanced interaction with Calnexin compared to ArcWT (Figure 7C-D).

## 4 Discussion

Previous work has established that disrupting the temporal dynamics of Arc by reducing ubiquitination leads to increased AMPA receptor endocytosis, enhanced mGluR-LTD, and impairments of spatial reversal learning (Wall et al., 2018); however, the intracellular mechanisms by which these phenotypes occurred were unknown. Here we found that disrupting Arc ubiquitination leads to altered ER Ca^2+^ release dynamics in all regions of the neuron except for secondary branchpoints. This phenotype was correlated with increased Arc-CaMKII interactions and Arc self-association. We also demonstrated that Arc ubiquitination and/or turnover may regulate its association with calnexin, providing a possible link between Arc ubiquitination levels and ER functionality.

As our previous work demonstrated that mGluR levels remained the same in ArcKR hippocampus relative to ArcWT (Wall et al., 2018), we hypothesized that dysregulation downstream of mGluR activation, such as ER Ca^2^ release, may be disrupted in ArcKR neurons. Indeed, we observed that ArcKR neurons release excess ER Ca^2+^ in response to DHPG. While Arc has previously been shown to interact with the cytoplasmic domain of the ER-associated protein calnexin via its central linker region (Myrum et al., 2017), to our knowledge there has never been a study directly implicating Arc in the modulation of ER Ca^2^ release dynamics. Interestingly, our coimmunoprecipitation data suggest an increased association of ArcKR with calnexin. However, it is unclear if this increased interaction would lead to excess ER Ca^2^ release in neurons. It is notable that the activity of SERCA, which is responsible for ER Ca^2+^ refilling (Berridge et al., 2003; Vandecaetsbeek et al., 2009), is modulated by and directly interacts with calnexin (Roderick et al., 2000; Lynes et al., 2013; Gutiérrez et al., 2020). Although we did not observe an interaction of ArcWT and ArcKR with SERCA, it is possible that a deficiency in Arc ubiquitination may indirectly reduce SERCA activity, perhaps by altering interactions between calnexin and SERCA. However, further work is necessary to test this hypothesis.

We observed that ArcKR has enhanced interactions with phosphomimetic forms of CaMKIIa and CaMKIIb. CaMKII is important for mGluR-LTD magnitude (Mockett et al., 2011) and its interaction with Arc has previously been established (Okuno et al., 2012; Zhang et al., 2019). Mutations in the CaMKIIa T286 autophosphorylation site were found to block DHPG-induced LTD in hippocampal slices (Zhang et al., 2019). Remarkably, ArcKR neurons exhibit increased GluA2 surface trafficking (Wall et al., 2018), and activated CaMKII has been shown to work in complex with another calcium-regulated protein PICK1 (protein interacting with C-kinase 1) to facilitate GluA2 trafficking from the ER to the cell surface in an ER Ca^2+^ release-dependent manner (Lu et al., 2014). Notably, GluA2 undergoes Q/R editing, decreasing AMPA receptor Ca^2+^ permeability (Bowie and Mayer, 1995; Plant et al., 2006; Zhou et al., 2017), hinting at a potential compensatory effect of elevated calcium signaling in ArcKR neurons. Arc itself is known to interact with PICK1 via the PICK1 BAR domain, with the interaction and spatial distribution of the two proteins altered under depolarizing conditions (Goo et al., 2018); whether this interaction is influenced by the ArcKR mutation is not known.

We found that ArcKR has increased self-association compared to ArcWT. Using SDS-PAGE under non-reducing conditions, we observed signs that ArcKR may be biased for increased oligomerization. Interestingly, Arc oligomerization is negatively regulated via phosphorylation at Ser 260 by CaMKII. Mutation of CaMKII phosphorylation sites on Arc leads to enhanced mGluR-LTD and causes impairments in the acquisition of fear induced memories (Zhang et al., 2019). Mirroring that work, we also found that ArcKR mice have impaired fear memories (data not shown). Given these correlates, one possibility is that the ArcKR mutation reduces CamKII-dependent phosphorylation at Ser 260. Conversely, it is unknown if Arc CamKII-dependent phosphorylation sites regulate its ubiquitination which may alter ER-mediated Ca^2+^ release.

Given the literature and our findings, we hypothesize a mechanism whereby CaMKII-dependent phosphorylation of Arc promotes Arc ubiquitination, which in turn tempers ER-mediated Ca^2+^ release to promote synaptic strengthening via reducing AMPA receptor endocytosis. In this model, CaMKII auto-phosphorylation marks a synapse for strengthening, and p-CaMKII interacts preferentially with non-ubiquitinated Arc. CamKII then phosphorylates Arc at Ser 260, increasing its ubiquitination. This serves to counteract Arc activity by 1) attenuating type 1 mGluR-activation-mediated ER Ca^2+^ release, 2) promoting Arc degradation via the ubiquitin-proteasome pathway, and 3) subsequently tempering AMPA receptor endocytosis. This mechanism would explain the increased ArcKR interaction with CaMKII. An additional consideration is that Arc ubiquitination also regulates Ser 260 phosphorylation. Given our observed spatial differences in Arc localization in neurons and correlations to ER-mediated Ca^2+^ release, this mechanism could be a way in which neurons provide regional specificity along dendritic regions to fine-tune synaptic strength in response to select synaptic inputs.

Arc-dependent intracellular Ca^2+^ dysregulation carries important implications for behavioral output. ArcKR mice exhibit impaired spatial reversal learning on a Barnes maze task which was associated with enhanced mGluR-LTD (Wall et al., 2018). Further work is aimed at determining whether this behavioral deficit is also due in part to other ArcKR effects observed in this study. For example, dysregulation of CaMKII phosphorylation has been shown to affect strategy selection on the Barnes maze (Bach et al., 1995). Furthermore, n-methyl-d-aspartate receptor-mediated LTD is required for behavioral flexibility on a similarly spatial learning-dependent Morris water maze task (Nicholls et al., 2008; Kim et al., 2011). Prevention of LTD induction and reduction of AMPA receptor endocytosis caused by stabilization of the adhesion molecule β-catenin causes similar deficits in reversal learning on the water maze (Mills et al., 2014). Enhanced Arc translation is found to occur in Fragile X-syndrome model mice, which also display enhanced mGluR-LTD and increased AMPA receptor endocytosis (Hou et al., 2006; Nakamoto et al., 2007; Cheng et al., 2017). It is possible that this model has increased DHPG-induced ER-Ca^2+^ release similar to our ArcKR mice. Tempering ER-Ca^2+^ release via increases in Arc ubiquitination might serve as an alternative mechanism that could restore excessive mGluR-LTD in this mouse model.

Cumulatively, our ArcKR model suggests that the cooperation of mGluR-LTD within an optimal magnitude, threshold, and temporal range are necessary for intracellular signaling events that control behavioral outputs - whether some of these are regulated by Arc ubiquitination, Arc-CaMKII interaction, and regulation of ER Ca^2+^ release is a subject for future research.

## 6 Author Contributions

MAG designed the study, performed experiments, and wrote the manuscript. ZDA, CLM, and BD performed experiments. NF and JJY contributed to study design. AMM designed the study, edited the manuscript, and contributed to manuscript writing.

## 8 Acknowledgments

We would like to thank Dr. Ulli Bayer for providing the CaMKII constructs, the Neuroscience Institute core staff, and GSU Division of Animal Resources.

Dr. Bin Dong is now affiliated with University of Arkansas, Department of Chemistry and Biochemistry, Fayetteville, Arkansas 72701. Dr. Ning Fang is now affiliated with Xiamen University, College of Chemistry and Chemical Engineering, Xiamen, China 361005.

